# GSK3α/β restrains IFNγ-inducible costimulatory molecule expression in alveolar macrophages, limiting CD4^+^ T cell activation

**DOI:** 10.1101/2023.08.16.553574

**Authors:** Laurisa M. Ankley, Kayla N. Conner, Taryn E. Vielma, Mahima Thapa, Andrew J Olive

**Affiliations:** Department of Microbiology and Molecular Genetics, College of Osteopathic Medicine, Michigan State University, East Lansing, MI 48824, USA

## Abstract

Macrophages play a crucial role in eliminating respiratory pathogens. Both pulmonary resident alveolar macrophages (AMs) and recruited macrophages contribute to detecting, responding to, and resolving infections in the lungs. Despite their distinct functions, it remains unclear how these macrophage subsets regulate their responses to infection, including how activation by the cytokine IFNγ is regulated. This shortcoming prevents the development of therapeutics that effectively target distinct lung macrophage populations without exacerbating inflammation. We aimed to better understand the transcriptional regulation of resting and IFNγ-activated cells using a new *ex vivo* model of AMs from mice, fetal liver-derived alveolar-like macrophages (FLAMs), and immortalized bone marrow-derived macrophages (iBMDMs). Our findings reveal that IFNγ robustly activates both macrophage types; however, the profile of activated IFNγ-stimulated genes varies greatly between these cell types. Notably, FLAMs show limited expression of costimulatory markers essential for T cell activation upon stimulation with only IFNγ. To understand cell type-specific differences, we examined how the inhibition of the regulatory kinases GSK3α/β alters the IFNγ response. GSK3α/β controlled distinct IFNγ responses, and in AM-like cells, we found GSK3α/β restrained the induction of type I IFN and TNF, thus preventing the robust expression of costimulatory molecules and limiting CD4^+^ T cell activation. Together, these data suggest that the capacity of AMs to respond to IFNγ is restricted in a GSK3α/β-dependent manner and that IFNγ responses differ across distinct macrophage populations. These findings lay the groundwork to identify new therapeutic targets that activate protective pulmonary responses without driving deleterious inflammation.

## INTRODUCTION

Macrophages are innate immune cells that play an important role in sensing the environment, initiating inflammation, and helping to activate the adaptive immune response (1). In the lungs, both resident and recruited macrophages play important roles in maintaining pulmonary function and protecting against respiratory pathogens (2). Resident lung macrophages, known as alveolar macrophages (AMs), reside in the airspace to recycle surfactants produced by the lungs (3). AMs are the first immune cells to detect pathogens in the lungs and are tasked with appropriately responding to stimuli while maintaining pulmonary function (4). During respiratory infections, monocyte-derived inflammatory macrophages are recruited to the lung tissues to support antimicrobial responses and resolve infections (5, 6). Dysregulation of these two important macrophage populations can result in pulmonary dysfunction, susceptibility to infection, and autoinflammatory disease (3, 7, 8). While both resident and recruited macrophages contribute to immune responses in the lungs, their regulation and functional mechanisms are distinct (9). For example, several studies suggest that AMs and recruited macrophages have different abilities to activate protective T cell responses yet how this process is regulated remains unkown (10–12).

One cue that drives T cell interactions with macrophages is the cytokine IFNγ. IFNγ binds to IFNγR on macrophages, activating Jak/Stat signaling pathways to drive the transcriptional induction of hundreds of genes that are mediated by interferon regulatory factors (IRFs) (13). Among these IFNγ-inducible genes are important T cell modulatory markers, including antigen presentation machinery, such as MHCI and MHCII, as well as costimulatory/inhibitory markers, including CD40, CD80, CD86, and PD-L1. In bone marrow-derived macrophages (BMDMs), IFNγ responses can be further fine-tuned through the activity of key regulators including the kinases GSK3α/β and mTOR (14–17). It remains to be understood whether this regulation is conserved in both AMs and recruited macrophages and how this contributes to differences in T cell activation.

Developing new therapies to combat respiratory infections will require a mechanistic knowledge of regulatory and functional differences between distinct macrophage subsets, to ensure effective control of inflammation and T cell effector function in the lungs. However, this understanding requires *ex vivo* models that faithfully recapitulate *in vivo* macrophage biology. BMDMs are differentiated from myeloid progenitors and are a widely used model for recruited inflammatory macrophages (18). Following activation with IFNγ, BMDMs become highly glycolytic, driving inflammatory cytokine production and directly modulating T cell responses in a similar manner to recruited macrophages (14, 19, 20).

Until recently, the availability of *ex vivo* models for AMs posed a challenge as AMs are notoriously difficult to maintain and isolate (21, 22). Thus, this technical hurdle has limited our understanding of regulatory networks that control AM functionality. It is important to understand how these lung-resident cells uniquely respond to inflammatory signals compared to other macrophages to develop lung-specific therapies that protect against infection while maintaining pulmonary function. To address this gap, several groups developed approaches to culture AM-like cells *ex vivo* that maintain AM functions (21, 23–25). While the details of these approaches differ, they all leverage lung-specific cytokine cues from GM-CSF and TGFβ that are required to maintain AM populations in the lung environment. We previously developed an *ex vivo* AM model known as fetal liver-derived alveolar-like macrophages (FLAMs) that uses fetal liver cells, which are AM progenitors, to interrogate AM function. Our previous work showed that FLAMs maintain high expression of the AM surface marker SiglecF and the key transcription factor Pparγ (21). FLAMs are easy to isolate, culture, and expand, which along with their genetic tractability, are strengths that allow mechanisms underlying AM functions to be interrogated.

Here, we examined the transcriptional profile of resting and IFNγ-activated FLAMs and immortalized BMDMs (iBMDMs) to better define functional differences between these key macrophage subsets. Our results show that FLAMs are similar to primary AMs, and while both FLAMs and iBMDMs respond to IFNγ, they exhibit unique transcriptional profiles. The regulation of these IFNγ responses is also distinct, with GSK3α/β playing unique roles in FLAMs and iBMDMs. Modulating GSK3α/β activity in IFNγ-activated FLAMs results in the robust production of type I IFN, which contributes to the induction of costimulatory molecules and increases the capacity of both FLAMs and AMs to activate CD4^+^ T cells. Our results suggest that AMs are restrained in their capacity to activate CD4^+^ T cells following IFNγ stimulation and that the IFNγ response is uniquely regulated in different macrophage subsets. These results have implications when considering host-directed therapies that target distinct macrophage populations in the pulmonary tissue.

## RESULTS

### FLAMs are phenotypically similar to AMs and distinct from BMDMs

To gain a global understanding of FLAM transcriptional patterns and to identify similarities and differences between FLAMs and other macrophage populations, we conducted RNA sequencing (RNA-seq) analysis of resting FLAMs and iBMDMs. Using differential expression analysis, we identified hundreds of genes that were significantly different between FLAMs and iBMDMs (Figure 1A and TableS1). These data suggest that FLAMs and iBMDMs are transcriptionally distinct macrophage populations.

**Figure 1.**
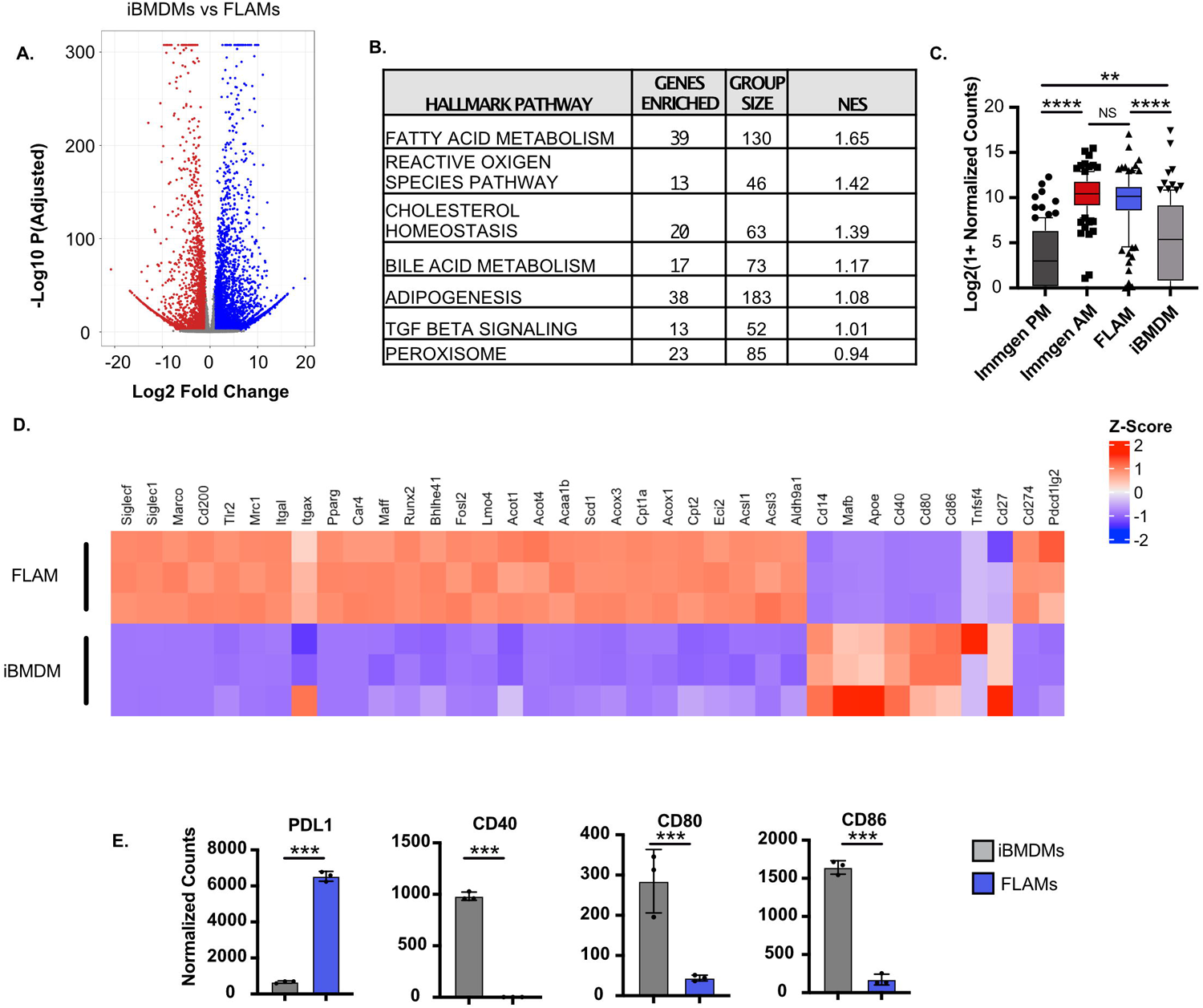
FLAMs are genetically similar to AMs. **(A)** Differentially expressed genes in untreated FLAMs (blue) and iBMDMs (red) were identified using RNA-seq. **(B)** Top 7 hallmark pathways enriched in untreated FLAMs. **(C)** Normalized counts of core AM genes were compared among iBMDMs, FLAMs, and previously published datasets (Immgen PM and Immgen AM). The box plot shows the median with quartiles representing the 10^th^–90^th^ percentile range of the data within that cell type. Each point represents the mean normalized counts of an individual gene. The Mann–Whitney U test was used to make statistical comparisons between each cell type and to compare medians. **(D)** Relative expression of genes previously associated with recruited macrophages was compared between FLAMs and iBMDMs and expressed as a heatmap Color scale represents Z-Score calculated from normalized read counts across samples for each gene. **(E)** Normalized counts of costimulatory molecules between untreated FLAMs (blue) and iBMDMs (grey). Adjusted p-values were determined using DeSeq2. *All data points represent one biological replicate +/- the standard deviation from one experiment. ****p<0.0001, ***p<0.001, **p<0.01, *p<0.05*.

To identify global pathways that were uniquely enriched in FLAMs, gene set enrichment analysis (GSEA) was performed using a ranked gene list generated from the differential expression analysis. Among the top Hallmark pathways enriched in FLAMs, we identified fatty acid metabolism, TGFβ signaling, cholesterol homeostasis, and peroxisome pathways (Figure 1B and TableS2). Given that AMs drive lipid metabolism in a fashion that is dependent on the transcription factor Pparγ, these data suggest the FLAM transcriptional profile is similar to that of primary AMs (26). To directly evaluate similarity to primary macrophages, we compared the FLAM and iBMDM RNA-seq transcriptional profiles with previously published datasets examining primary AMs and peritoneal macrophages (27). In line with the GSEA results, we found that FLAMs were more similar to AMs, whereas iBMDMs were more similar to peritoneal macrophages (Figure 1C and Table S3). These findings suggest that FLAMs are a robust *ex vivo* model for AMs.

To further examine similarities with primary macrophages, we studied a subset of genes that were previously associated with recruited macrophages or AMs (28). We found iBMDMs expressed high levels of genes associated with recruited macrophages, including CD14, ApoE, and the key transcription factor MafB (Figure 1D and TableS4). In contrast, FLAMs expressed high levels of transcription factors associated with resident lung macrophages, such as Pparγ, Car4, Maff, Fosl2, Bhlhe41, and Runx2. In addition, FLAMs expressed high levels of resident macrophage associated surface markers, including SiglecF, Siglec1, Marco, CD200, TLR2, MRC1, Itgal, and Itgax, which were expressed at low levels or not expressed in iBMDMs. In line with there being functional similarities between AMs and FLAMs, we observed high expression of genes associated with lipid and cholesterol metabolism in FLAMs (29). Interestingly, when we examined genes that modulate T cell activation, we observed high expression of the coinhibitory markers PDL1 and PDL2 on FLAMs (Figure 1D and 1E) (30). In contrast, we observed very low expression of costimulatory molecules, including CD40, CD80, and CD86 (Figure 1D and 1E). These data show that FLAMs express core AM-associated genes, unlike iBMDMs.

To confirm our transcriptional results with an orthologous method, we compared the expression of surface markers predicted to be differentially expressed on AMs and FLAMs relative to iBMDMs with flow cytometry. We found the surface markers CD11a, TLR2, MRC1, and Siglec1 were all highly expressed on both resting FLAMs and primary AMs, while resting iBMDMs expressed higher levels of CD14 (Figure 2A and 2B). In agreement with our transcriptional profiling, we also found low expression of costimulatory markers on FLAMs and AMs compared to iBMDMs but high expression of the coinhibitory marker PD-L1 on FLAMs and AMs (Figure 2C and 2D). Taken together, these results show that FLAMs are transcriptionally distinct from iBMDMs and are a good model for primary AMs. Additionally, in resting conditions, FLAMs and AMs express low levels of T cell-activating costimulatory markers.

**Figure 2.**
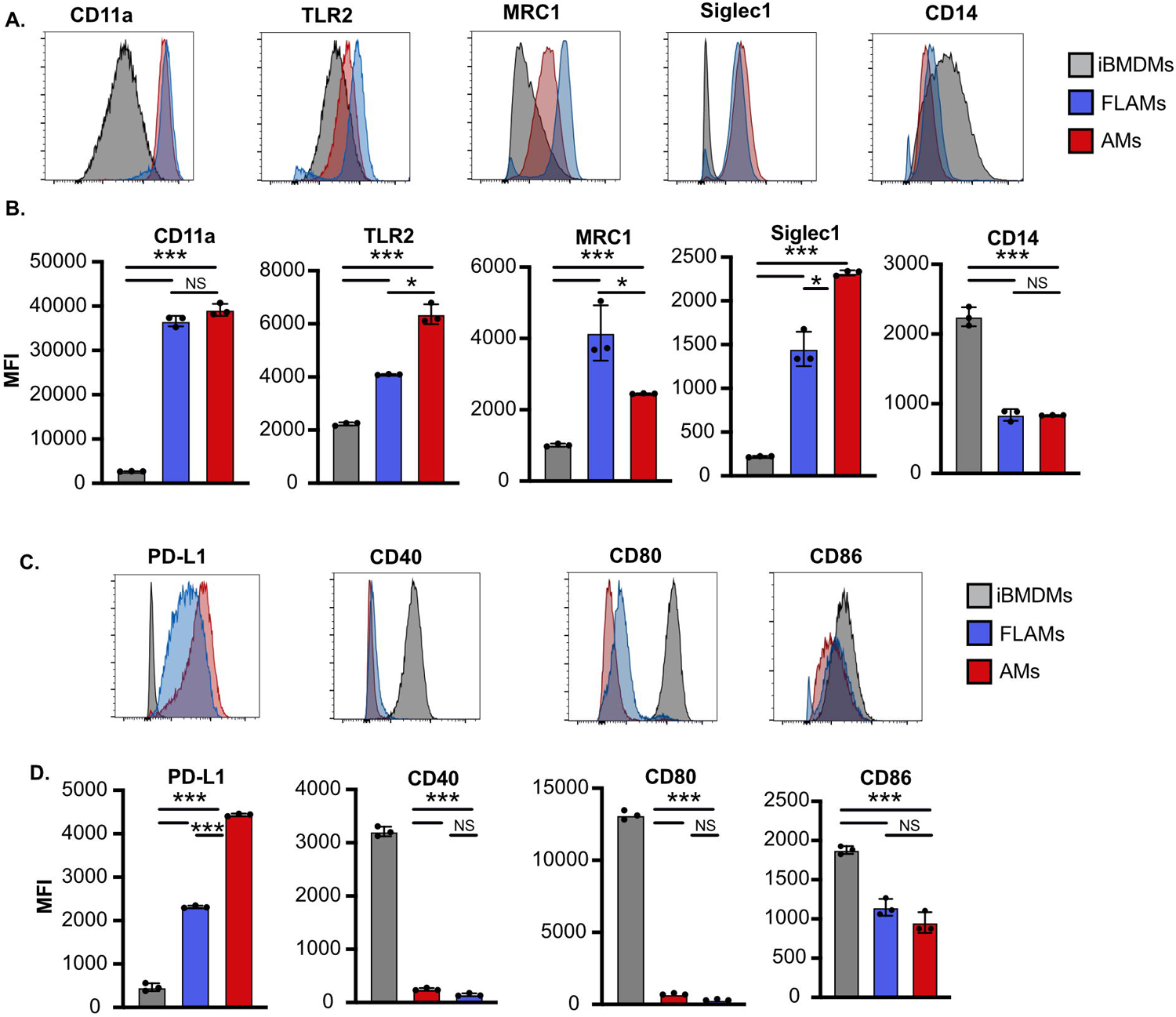
FLAMs express surface markers associated with AMs. **(A)** Representative histograms were overlaid for selected surface markers expressed on AMs (red), FLAMs (blue), or iBMDMs (grey). **(B)** The mean fluorescence intensity (MFI) of each surface marker on each cell type was quantified. One-way ANOVA with a Tukey test for multiple comparisons were used. **(C)** Representative histograms were overlaid for selected costimulatory markers expressed on AMs (red), FLAMs (blue), or iBMDMs (grey). **(D)** The mean fluorescence intensity (MFI) of each costimulatory marker was quantified. One-way ANOVA with a Tukey test for multiple comparisons were used. *All data points represent one biological replicate +/- the standard deviation from one experiment of three. ****p<0.0001, ***p<0.001, **p<0.01, *p<0.05*.

### IFN**γ** induces distinct transcriptional profiles in FLAMs and does not broadly induce T cell costimulatory molecules in FLAMS

The cytokine IFNγ acts as an important regulator of the host response in macrophages by inducing the expression of antimicrobial molecules and T cell modulatory molecules to help drive protective immune responses (31–33). Since transcriptional differences are observed between FLAMs and iBMDMs at baseline, we wondered whether IFNγ responses would be similar or distinct in these two cell types. To address this question, we conducted global RNA-seq analysis of FLAMs and iBMDMs following IFNγ activation for 24 hours. We first used differential expression analysis to compare IFNγ-activated FLAMs or iBMDMs to their resting counterparts that were described above. For both iBMDMs and FLAMs, IFNγ stimulation resulted in the induction of hundreds of genes (Figure 3A, 3B and TableS1). This finding suggests that IFNγ robustly activates both iBMDMs and FLAMs.

**Figure 3.**
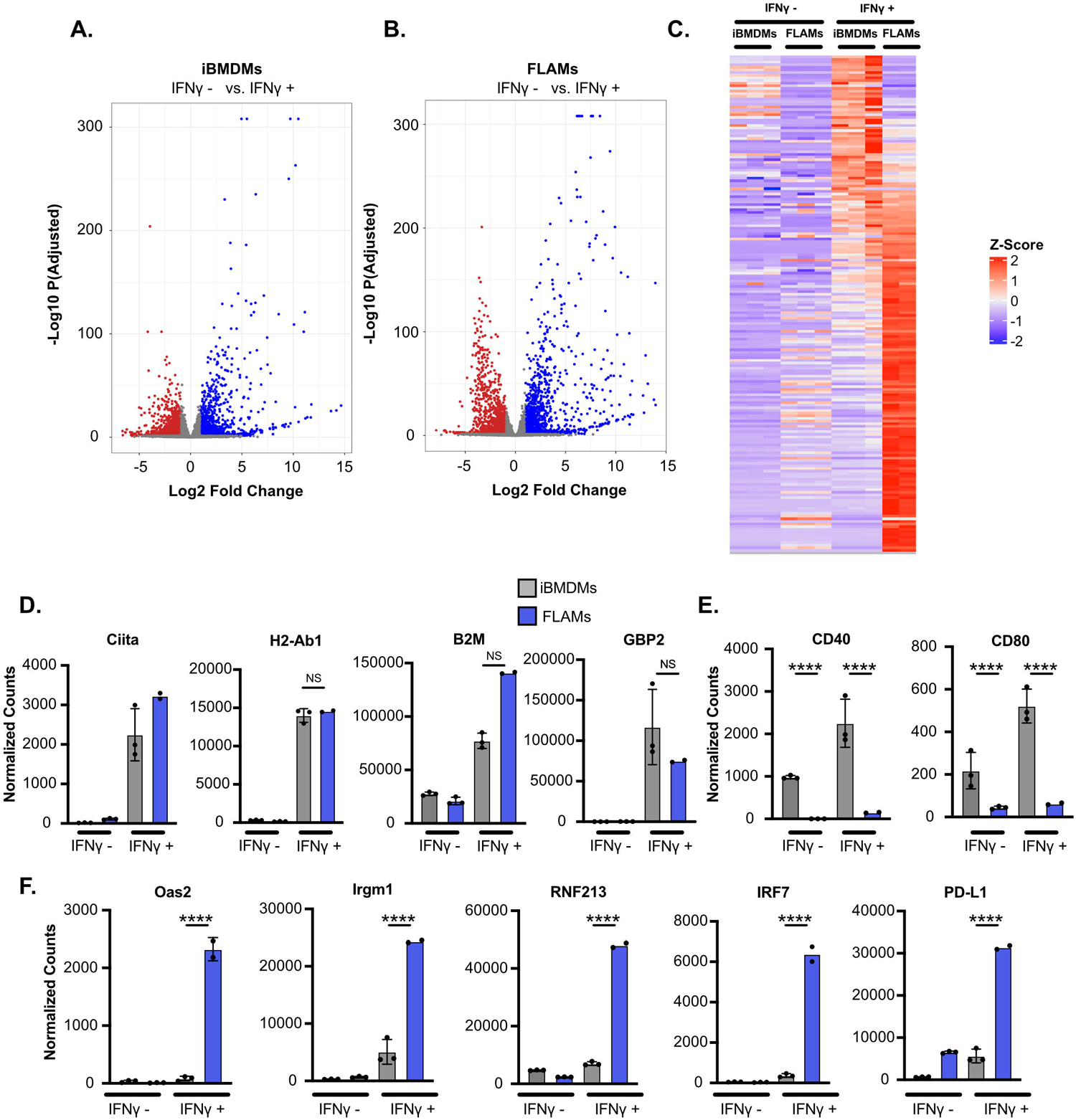
iBMDMs and FLAMs respond differently to IFNγ stimulation. **(A)** Differential expression of untreated (red) and IFNγ-stimulated (6.25 ng/mL) (blue) iBMDMs; colored symbols have an adjusted p-value <0.05 and a fold change greater than 2. **(B)** Differential expression of untreated (red) and IFNγ-stimulated (blue) FLAMs. Colored symbols have an adjusted p-value <0.05 and a fold change greater than 2. **(C)** Expression of a subset of ISGs was compared in untreated and IFNγ-stimulated iBMDMs and FLAMs and expressed as a heatmap Color scale represents Z-Score calculated from normalized read counts across samples for each gene. **(D)** Normalized counts of ISGs that were differentially regulated in iBMDMs (grey) and FLAMs (blue) with and without IFNγ. Statistical significance was determined based on adjusted p-values using DeSeq2. Data points represent one biological replicate from one experiment. **(E)** Normalized counts of costimulatory molecules that were differentially regulated in iBMDMs (grey) and FLAMs (blue) in untreated and IFNγ-stimulated conditions. **(F)** Normalized counts of cell autonomous restriction factors that were differentially regulated in iBMDMs (grey) and FLAMs (blue) in untreated and IFNγ-stimulated conditions. *Normalized count datapoints represent one biological replicate from one experiment +/- the standard deviation. MFI datapoints represent one biological replicate +/- the standard deviation from one representative experiment of three. ****p<0.0001, ***p<0.001, **p<0.01, *p<0.05*.

To directly compare the IFNγ-mediated responses of iBMDMs and FLAMs, we visualized the normalized reads for genes associated with a curated IFNγ-stimulated gene (ISG) set based on the hallmark pathway for IFNγ-activated cells (Figure 3C and Table S4). Antigen presentation machinery for MHCI and MHCII was robustly induced following activation of both FLAMs and iBMDMs (Figure 3D). However, not all ISGs were differentially induced in stimulated FLAMs and iBMDMs. For example, the costimulatory molecules CD40 and CD80 were robustly induced in iBMDMs, but their expression remained low in FLAMs (Figure 3E). In contrast, we noted cell-autonomous restriction factors, including OAS2, Irgm1, and RNF213, were induced at levels over 10-fold higher in FLAMS than iBMDMs (Figure 3F). Additionally, the transcription factor IRF7 was induced at levels 2-4-fold higher than baseline in iBMDMs following IFNγ activation, whereas in FLAMs, the observed induction was over 100-fold higher than at baseline. In line with our observations in resting cells, we found that the expression of the coinhibitory marker PDL1 remained over 10-fold higher following IFNγ activation in FLAMs than in iBMDMs. Taken together, these results show that while both FLAMs and iBMDMs robustly respond to IFNγ activation, this activation induces distinct transcriptional changes in both cell types, including differences in T cell costimulatory molecules that remained expressed at low levels in FLAMs.

### GSK3α/**β** inhibition during IFN**γ** activation of AMs and FLAMs results in the robust upregulation of costimulatory molecules

We next examined how the different IFNγ responses in FLAMs and iBMDMs are regulated. Previous work showed that GSK3α and GSK3β are key regulators that fine-tune the IFNγ response in iBMDMs and that inhibiting GSK3α/β in iBMDMs blocks a subset of IFNγ responses, including the expression of the MHCII transactivator Ciita and subsequent MHCII expression (14). However, the core IFNγ signaling pathways, including Stat1 and Irf1, remained intact following GSK3α/β inhibition. We hypothesized that GSK3α/β may contribute to the different IFNγ responses observed between iBMDMs and AMs. To test this hypothesis, we cultured resting or IFNγ-activated iBMDMs or FLAMs with and without the highly specific GSK3α/β inhibitor CHIR99021 and then analyzed MHCII expression with flow cytometry (34). In agreement with our previous results, GSK3α/β blockade in iBMDMs led to a significant reduction in MHCII on IFNγ-activated cells (Figure 4A). In contrast, inhibiting GSK3α/β in IFNγ-activated FLAMs increased MHCII expression (Figure 4A). These data suggest that GSK3α/β have distinct functions in controlling the IFNγ response in FLAMs and BMDMs.

**Figure 4.**
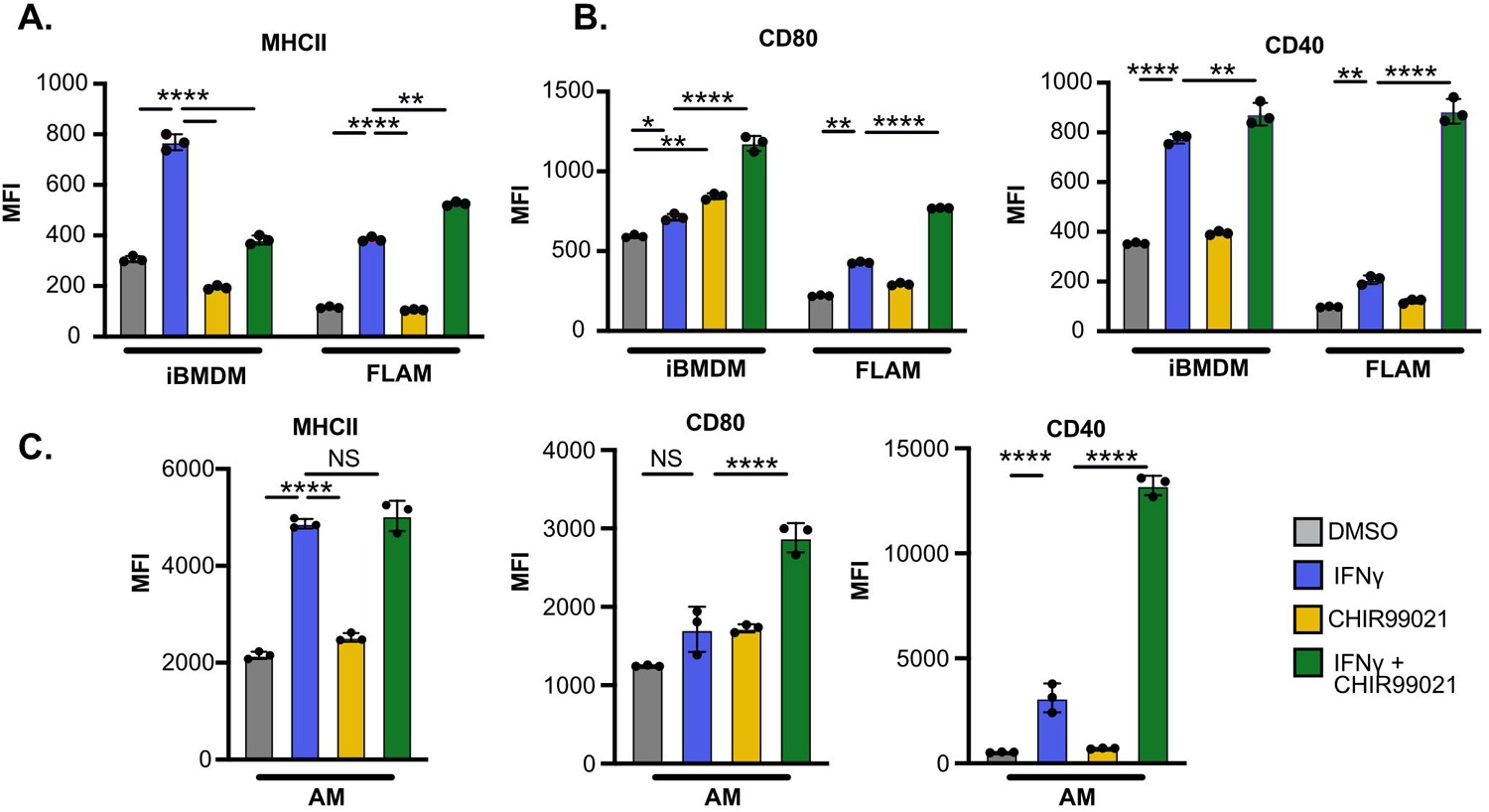
Blocking GSK3α/β during IFNγ activation of FLAMs drives costimulatory marker expression. iBMDMs and FLAMs treated with DMSO (grey), DMSO and IFNγ (6.25 ng/mL) (blue), CHIR (10 μM) (yellow), or CHIR and IFNγ (10 μM and 6.25 ng/mL) (green) for 24 hours. (**A)** The surface expression of MHCII and **(B)** the indicated costimulatory molecules was quantified by flow cytometry. Each datapoint represents the MFI for each biological replicate +/- the standard deviation from one representative experiment of four similar experiments. Statistical significance was determined with two-way ANOVA and a Tukey test for multiple comparisons. **(C)** Primary AMs were treated with DMSO (grey), DMSO and IFNγ (6.25 ng/mL) (blue), CHIR (10 μM) (yellow), or CHIR and IFNγ (10 μM and 6.25 ng/mL) (green) for 24 hours. The surface expression of the indicated T cell modulatory molecules was quantified by flow cytometry. Statistical significance was determined with two-way ANOVA and a Tukey test for multiple comparisons. *MFI datapoints represent one biological replicate +/- the standard deviation from one representative experiment of three. ****p<0.0001, ***p<0.001, **p<0.01, *p<0.05*.

GSK3α/β were previously shown to modulate costimulatory molecule expression in different cell types (35, 36). Thus, we next tested if GSK3α/β inhibition alters the IFNγ-mediated induction of costimulatory molecules. Resting or IFNγ-activated iBMDMs and FLAMs were treated with DMSO or CHIR99021, and flow cytometry was used to quantify the surface expression of CD40 and CD80. We found that while IFNγ increased the expression of all markers on iBMDMs, GSK3α/β blockade had no effect on this induction (Figure 4B). In contrast, while IFNγ alone resulted in minimal changes to costimulatory molecule expression on FLAMs, GSK3α/β blockade in IFNγ-activated FLAMs resulted in a robust increase in all costimulatory molecules. These results for MHCII and costimulatory markers were confirmed in primary AMs (Figure 4C). Together, these findings suggest that GSK3α/β play distinct functions in regulating the response to IFNγ in AMs and BMDMs.

### GSK3α/**β** inhibition during IFN**γ** activation robustly alters the transcriptional landscape of FLAMs

As GSK3α/β inhibition had different impacts on a subset of IFNγ responses in BMDMs and FLAMs, we next examined global transcriptional changes that occurred during GSK3α/β inhibition. RNA-seq analysis was conducted on resting and IFNγ-activated iBMDMs and FLAMs in the presence of CHIR99021, and these results were compared to the above RNA-seq analysis in resting and IFNγ-activated iBMDMs and FLAMs. First, we examined MHCII and costimulatory marker expression and found that in line with our flow cytometry results, FLAMS treated with combination IFNγ and CHIR99021 exhibited no change in H2-Ab1 (MHCII) expression but over 100-fold induction of CD80 and CD40 that was not observed in any other condition (Figure 5A and Table S1). Principal component analysis (PCA) of all 8 RNA-seq conditions (+/- IFNγ +/- CHIR99021 in iBMDMs and FLAMs) revealed clear differences in the transcriptional landscape of iBMDMs and FLAMs (Figure 5B). All iBMDM samples clustered closely within the PCA plot, with distinct but small shifts in the transcriptomes following IFNγ and/or GSK3α/β inhibition. Compared to resting iBMDMs, resting FLAMs were significantly shifted along PC1, in line with the above results showing distinct transcriptional landscapes in these resting cell types. While the shifts in the transcriptional profile of FLAMS in response to either IFNγ activation or GSK3α/β blockade were similar to those observed in iBMDMs, the combination of IFNγ and CHIR99021 resulted in a major shift in the transcriptional landscape of FLAMs along PC2. These results show that GSK3α/β are key regulators of the IFNγ response in FLAMs and that the combination of IFNγ activation and GSK3α/β blockade drives a synergistic transcriptional response not observed in any other FLAM or iBMDM condition.

**Figure 5.**
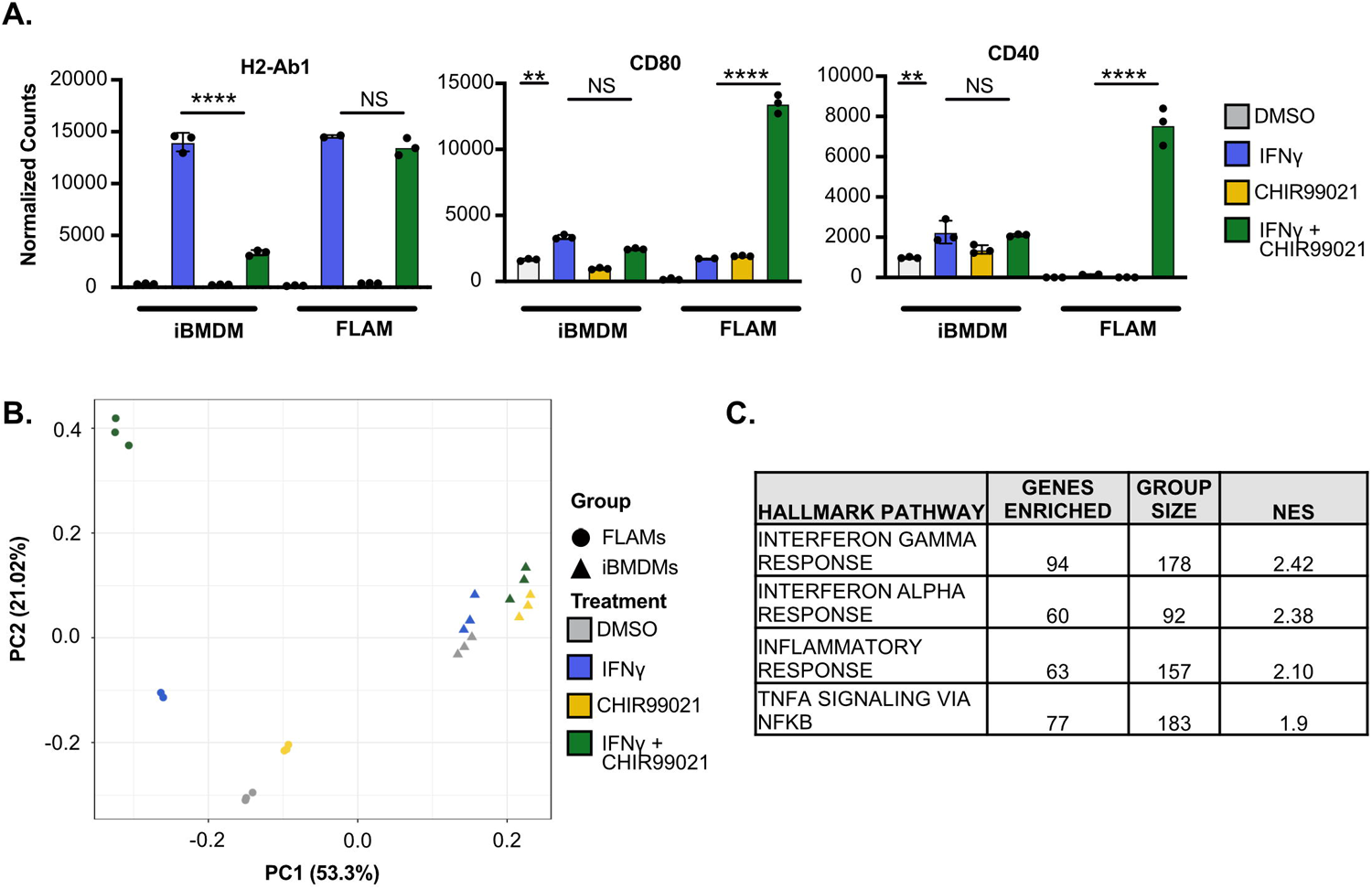
Inhibition of GSK3α/b in IFNγ-activated FLAMs changes the transcriptional landscape. iBMDMs and FLAMs were treated with DMSO (grey), DMSO and IFNγ (6.25 ng/mL) (blue), CHIR (10 μM) (yellow), or CHIR and IFNγ (10 μM and 6.25 ng/mL) (green) for 24 hours, and RNA-seq was completed. **(A)** Normalized counts of T cell activation molecules in iBMDMs and FLAMs are shown from RNA-seq analysis in the indicated conditions. Statistical significance was determined with adjusted p-values calculated by DeSeq2. **(E)** A PCA plot comparing the similarity of iBMDMs and FLAMs in the indicated conditions is shown. **(F)** The top 4 Hallmark pathways enriched using GSEA are shown to compare GSK3α/β-inhibited/IFNγ-stimulated FLAMs and IFNγ-stimulated FLAMs from a ranked list calculated by DeSeq2. *Normalized counts datapoints represent one biological replicate +/- the standard deviation from one experiment. MFI points represent one biological replicate +/- the standard deviation from one representative experiment of three. ****p<0.0001, ***p<0.001, **p<0.01, *p<0.05*.

To understand which pathways are altered during GSK3α/β inhibition in IFNγ-activated FLAMs, we next used GSEA based on a differential expression ranked list to compare IFNγ-activated FLAMs in the presence and absence of GSK3α/β inhibition. We found both IFNα and TNF pathways, in addition to IFNγ, were all significantly enriched in GSK3α/β-inhibited, IFNγ-activated FLAMs (Figure 5C and Table S2). These results suggest that blockade of GSK3α/β during IFNγ activation of FLAMs drives inflammatory cytokine responses that may contribute to the expression of key IFNγ-inducible genes, including costimulatory markers.

### Type I IFN and TNF contribute to the upregulation of costimulatory molecules on IFN**γ**-activated FLAMs when GSK3α/**β**are inhibited

We next interrogated the mechanisms driving costimulatory marker induction on GSK3α/β-inhibited, IFNγ-activated FLAMs. Our GSEA results identified TNF and IFNβ, which were previously associated with modulating costimulatory marker expression (20, 37). With our RNA-seq dataset, we found that in iBMDMs, TNF was expressed following IFNγ activation regardless of GSK3α/β inhibition; however, in FLAMs, TNF was highly expressed only following IFNγ activation and GSK3α/β inhibition (Figure 6A). IFNβ was not expressed in iBMDMs in any conditions, and high expression of IFNβ was observed only in IFNγ-activated, GSK3α/β-inhibited FLAMs. To confirm the results from RNA-seq analysis, we examined the production of cytokines using a multiplex Luminex assay of supernatants from resting and IFNγ-activated iBMDMs and FLAMs with and without GSK3α/β inhibition. In agreement with the transcriptomic studies, TNF and type I IFN were increased only in FLAMs following IFNγ-activation and GSK3α/β inhibition (Figure 6B). These data show that inhibition of GSK3α/β in IFNγ-activated FLAMs results in increased expression of costimulatory molecule-modulating cytokines.

**Figure 6.**
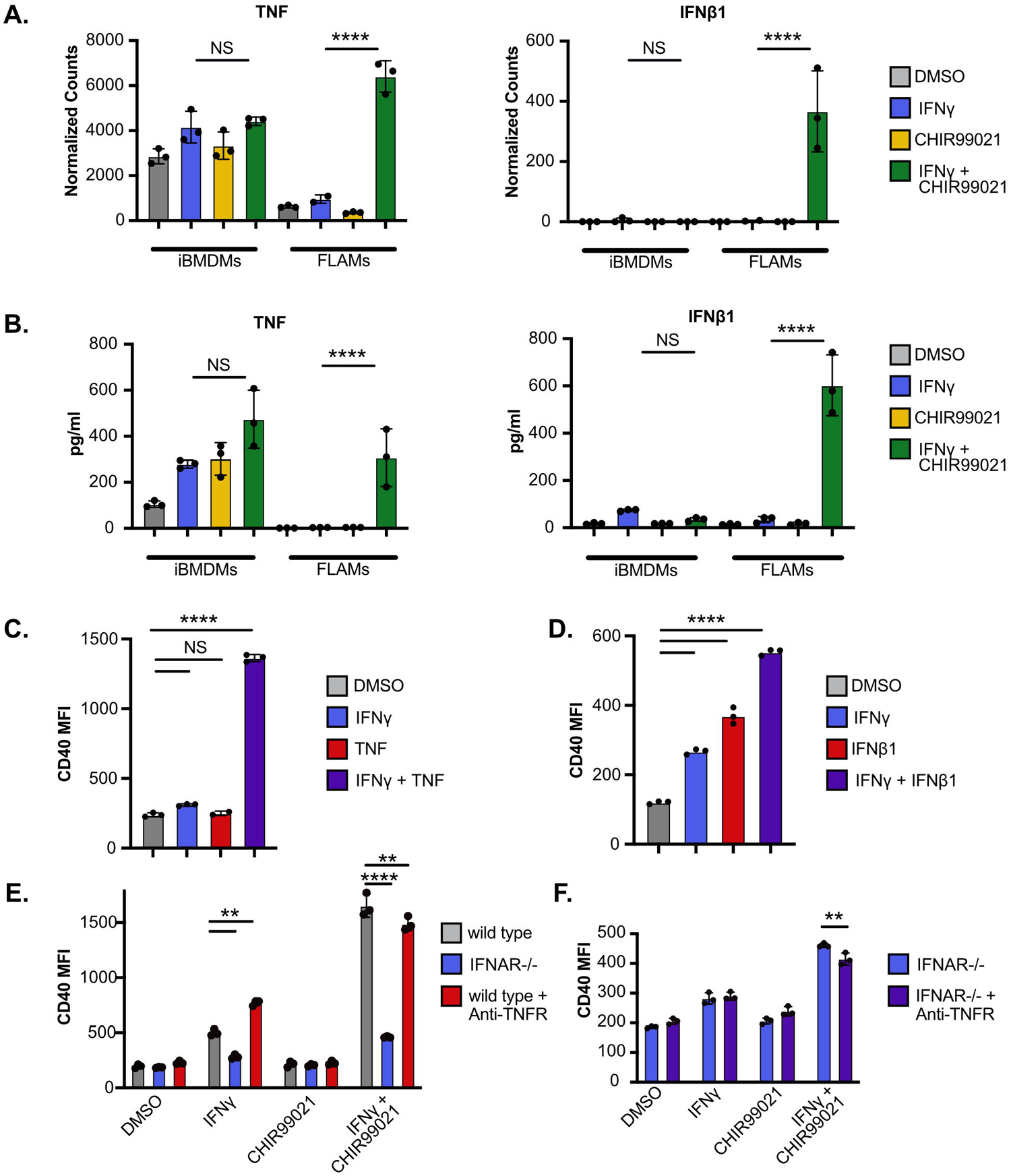
TNF and type I IFN contribute to CD40 expression on IFNγ-stimulated FLAMs. **(A)** Normalized counts are shown for TNF (left) and IFNβ (right) from iBMDMs and FLAMs treated with DMSO (grey), DMSO and IFNγ (6.25 ng/mL) (blue), CHIR (10 μM) (yellow), or CHIR and IFNγ (10 μM and 6.25 ng/mL) (green) for 24 hours. Statistical significance was determined with adjusted p-values using DeSeq2. **(B)** The production of TNF (left) and IFNβ (right) from iBMDMs and FLAMs in identical conditions as in A. Statistical significance was determined with two-way ANOVA and a Tukey test for multiple comparisons. **(C)** The MFI of CD40 for FLAMs treated with DMSO (grey), IFNγ (6.25 ng/mL) (blue), TNF (20 ng/mL) (red), or IFNγ and TNF (6.25 ng/mL and 20 ng/mL) (purple) for 24 hours. Statistical significance was determined with two-way ANOVA and a Tukey test for multiple comparisons. **(D)** The MFI of CD40 for FLAMs treated with DMSO (grey), IFNγ (6.25 ng/mL) (blue), IFNβ1 (20 ng/mL) (red), or IFNγ and IFNβ1 (6.25 ng/mL and 20 ng/mL) (purple) for 24 hours. Statistical significance was determined with two-way ANOVA and a Tukey test for multiple comparisons. **(E)** MFI of CD40 for WT (grey), IFNAR^-/-^ (blue), and WT + anti-TNF (red). FLAMs were cultured with the indicated treatments for 24 hours. Statistical significance was determined with two-way ANOVA and a Tukey test for multiple comparisons. **(F)** MFI of CD40 for IFNAR^-/-^ (blue) and IFNAR^-/-^ + anti-TNFR (purple) in the indicated conditions. Statistical significance was determined with one-way ANOVA and a Tukey test for multiple comparisons. *Normalized counts points represent three biological replicates +/- the standard deviation from one experiment. MFI and cytokine quantification points represent one biological replicate +/- the standard deviation from one representative experiment of three. ****p<0.0001, ***p<0.001, **p<0.01, *p<0.05*.

We next tested the sufficiency of either TNF or IFNβ to drive costimulatory marker expression on IFNγ-activated FLAMs. Resting or IFNγ-activated FLAMs were treated with recombinant TNF or IFNβ, and surface levels of CD40 were quantified with flow cytometry. While TNF alone did not increase CD40 expression on resting FLAMs, treatment of IFNγ-activated FLAMs with TNF resulted in a synergistic increase in CD40 expression (Figure 6C). The addition of type I IFN significantly increased CD40 expression in all conditions, and combination treatment with IFNγ and IFNβ resulted in higher CD40 expression than treatment with IFNβ alone (Figure 6D). These data suggest that both IFNβ and TNF contribute to the increased CD40 expression during IFNγ activation of FLAMs when GSK3α/β are inhibited.

Next, we tested whether the production of either TNF or IFNβ was required for enhanced costimulatory marker expression on GSK3α/β-inhibited, IFNγ-activated FLAMs. To block the function of IFNβ and TNF, we isolated FLAMs from IFNAR^-/-^ mice and used a TNFR neutralizing antibody, enabling the role of both cytokines to be tested simultaneously. Resting and IFNγ-activated wild-type and IFNAR^-/-^ FLAMs in the presence and absence of CHIR99021 and/or anti-TNFR antibodies were analyzed for CD40 expression by flow cytometry. We observed that TNF signaling blockade led to a minimal decrease in CD40 expression in IFNγ-activated, GSK3α/β-inhibited FLAMs, while IFNβ signaling blockade dramatically reduced CD40 expression (Figure 6E). When TNF was blocked in IFNAR^-/-^ FLAMs, a further reduction in CD40 surface expression was observed, although this change was relatively small. Taken together, these data suggest that both TNF and IFNβ contribute to the increase in costimulatory marker expression seen in IFNγ-activated, GSK3α/β-inhibited FLAMs.

### Inhibition of GSK3α/**β**following IFN**γ** activation of FLAMs and AMs drives CD4^+^ T cell activation

Costimulatory marker expression is necessary to activate the adaptive immune response during infection (38). Our previous studies found that the increase in antigen presentation and costimulatory markers following IFNγ activation of BMDMs is sufficient to activate CD4^+^ T cells (14, 20). Given that costimulatory marker expression was not induced in AMs or FLAMs with IFNγ alone but only in combination with GSK3α/β inhibition, we hypothesized that IFNγ-activated AMs or FLAMs would not robustly activate CD4+ T cells, whereas IFNγ-activated, GSK3α/β-inhibited cells would. To test this hypothesis, we used a previously optimized coculture assay with macrophages and TCR-transgenic CD4^+^ T cells that are specific for the *Mycobacterium tuberculosis* peptide p25 and assessed T cell activation in various conditions based on the production of IFNγ (39–41). We found that p25 CD4^+^ T cells alone produced no IFNγ, while coculture with peptide-pulsed splenocytes resulted in robust IFNγ production (Figure 7A). As expected, p25 CD4^+^ T cells cocultured with resting or CHIR99021-treated iBMDMs or FLAMs did not produce IFNγ. p25 CD4^+^ T cells cocultured with IFNγ-activated iBMDMs produced IFNγ, while GSK3α/β blockade in IFNγ-activated iBMDMs prevented p25 CD4^+^ T cell activation. In FLAMs, IFNγ activation alone was insufficient to activate p25 CD4^+^ T cells during coculture. In contrast, GSK3α/β inhibition in IFNγ-activated FLAMs resulted in the robust production of IFNγ by p25 CD4^+^ T cells. Similar results were observed when this experiment was repeated with primary AMs (Figure 7B). p25 CD4^+^ T cells were only activated upon coculture with AMs that were IFNγ-activated with GSK3α/β inhibition. Taken together, these data suggest that distinct macrophage populations have different abilities to directly activate T cell responses and highlight that AMs are restrained in their capacity to directly activate adaptive immune responses.

**Figure 7.**
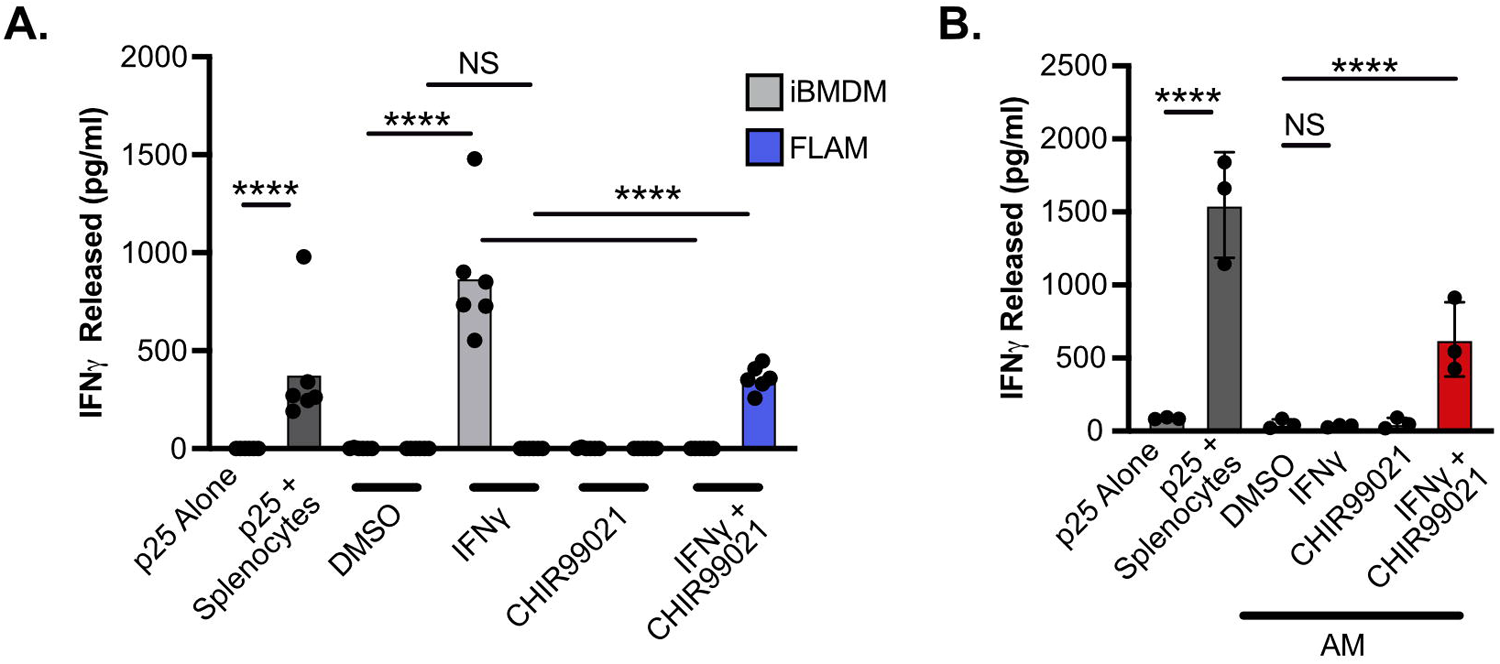
GSK3α/b restrain the ability of AMs to activate CD4^+^ T cells following IFNγ activation. **(A)** p25 peptide-pulsed iBMDMs and FLAMs treated with DMSO (grey), DMSO and IFNγ (6.25 ng/mL) (blue), CHIR (10 μM) (yellow), or CHIR and IFNγ (10 μM and 6.25 ng/mL) (green) for 24 hours were cocultured with naive p25-specific T cells for 4 days, and IFNγ released into the supernatant was quantified by ELISA. P25 T cells alone or P25 T cells with peptide-pulsed splenocytes were used as positive and negative controls for the assay. **(B)** p25 peptide-pulsed AMs treated with DMSO (grey), DMSO and IFNγ (6.25 ng/mL) (blue), CHIR (10 μM) (yellow), or CHIR and IFNγ (10 μM and 6.25 ng/mL) (green) for 24 hours were cocultured with naive p25-specific T cells for 4 days, and IFNγ released into the supernatant was quantified by ELISA. Statistical significance was determined with two-way ANOVA and a Tukey test for multiple comparisons. *Cytokine quantification points represent one biological replicate +/- the standard deviation from one representative experiment of two. ****p<0.0001, ***p<0.001, **p<0.01, *p<0.05*.

## DISCUSSION

While AMs are essential for lung function, experimental limitations have prevented a mechanistic understanding of how they uniquely respond to inflammatory signals. Here, we used FLAMs as an *ex vivo* model of AMs to globally understand transcriptional and functional differences between resident and recruited lung macrophages (21). By comparing the transcriptome of iBMDMs and FLAMs to previously published datasets from Immgen on various myeloid-derived cells, we found strong similarity between FLAMs and AMs but not between AMs and BMDMs(27). Examination of pathways associated with FLAMs and AMs identified signatures previously associated with AMs, including activation of Ppar signaling, unsaturated fatty acid synthesis, lipid metabolism, and lysosome and peroxisome pathways (29, 42–46). We also confirmed the expression of AM-associated surface markers including CD11a and Siglec1 in FLAMs, which were expressed at low levels on iBMDMs. Thus, we have convincingly shown that FLAMs are a tractable model that can be leveraged to dissect mechanisms regulating AM maintenance and function.

We used FLAMs to dissect how lung macrophages respond to the proinflammatory cytokine IFNγ. IFNγ is an important regulator of immunity in the lungs and activates a range of ISGs that drive adaptive immunity, cell autonomous effectors, and cytokines/chemokines (13, 47). Interestingly, our transcriptional profiling found that while both iBMDMs and FLAMs are responsive to IFNγ stimulation, they induce distinct transcriptional changes following activation. Given the metabolic differences between FLAMs and iBMDMs, our current model suggests that baseline metabolism and differences in IFNγ-mediated shifts in metabolism drive distinct responses to IFNγ. Previous studies in BMDMs showed that IFNγ activation drives a shift in cells towards aerobic glycolysis that is dependent on the activation of HIF1α (48). However, both aerobic glycolysis and oxidative phosphorylation are known to contribute to IFNγ responses (20, 49). Whether HIF1α plays a role in the different IFNγ responses between BMDMs and AMs and how metabolism shifts in AMs following IFNγ activation will be directly examined in the future. By coupling genetic approaches to remove single transcription factors with metabolic flux approaches including Seahorse assays, we will be well positioned to understand the mechanisms driving the interlinked metabolic and transcriptional responses following AM activation with IFNγ.

The metabolism-associated kinases GSK3α/β were previously associated with macrophage functions including IFNγ activation in iBMDMs (50, 51)(52). Thus, we examined the role of GSK3α/β in FLAMs. In contrast to iBMDMs, we found that GSK3α/β do not control IFNγ-dependent MHCII upregulation in FLAMs or AMs, and our transcriptional profiling showed that there are macrophage subset-specific roles for GSK3α/β in regulating the IFNγ response. Unlike in iBMDMs, the inhibition of GSK3α/β in combination with IFNγ activation in FLAMs resulted in a dramatic shift in the transcriptional landscape. It was beyond what was seen with either GSK3α/β inhibition or IFNγ activation alone. What drives the synergistic response of AMs to both IFNγ and GSK3α/β inhibition remains an open question. One potential explanation for this synergy is the observation that type I IFN responses are robustly induced only in IFNγ-activated FLAMs with GSK3 inhibition. Type I IFNs can be induced by endogenous ligands from the mitochondria, such as mitochondrial DNA or RNA, as well as changes in cholesterol metabolism (53–55). The contribution of these distinct IFN pathways in IFNγ-activated GSK3α/β-inhibited FLAMs and whether type I IFNs drive the observed transcriptional changes will need to be tested in the future. Altogether, our results show that GSK3α/β are important regulators of IFNγ responses in AMs that differ from those in BMDMs.

Throughout our study, we noticed major differences in the regulation of T cell modulatory markers between iBMDMs and FLAMs. While costimulatory molecules were robustly induced in iBMDMs with IFNγ activation alone, these molecules were only induced in FLAMs if GSK3α/β were also inhibited. These differences in MHCII and costimulatory molecule expression had functional implications, as FLAMs and AMs only activated CD4^+^ T cells during IFNγ activation if GSK3 was also blocked. Our data support different roles for various macrophage subtypes in directly modulating the adaptive immune system. Previous studies suggest that AMs are not efficient activators of naïve T cells (10, 11). In fact, robust activation of T cells by AMs was associated with worse clinical outcomes during infection with SARS-CoV-2 in a manner that was dependent on both IFNγ and TNF (33). We speculate that AMs respond to IFNγ in a way that prevents overly robust activation of T cells and limits deleterious lung damage. However, when combined with other inflammatory signals, including type I IFN or TNF, IFNγ drives AMs to robustly activate T cell responses. It is possible that pathogens such as *Mycobacterium tuberculosis* take advantage of the restrained T cell-activating capacity of AMs to prevent detection and initiate lung infections, but this hypothesis needs to be directly tested.

The strength of the approach presented here lies in the ease and tractability of using FLAMs as a model of AMs to identify mechanisms that make AMs unique. Throughout our study, significant differences were observed between iBMDMs and FLAMs in gene expression, surface protein expression, innate immune pathway activation, and the functional capacity to activate CD4^+^ T cells. Our data suggest an urgent need to define AM-specific regulation of host responses rather than extrapolating immune pathway regulatory mechanisms previously observed in BMDMs. Filling this gap may identify potential targets for lung-specific therapies and provide a better understanding of how AMs initiate immune responses while maintaining pulmonary function. It will also be important to dissect these pathways directly within the lung environment, which is a limitation of our current study. Future work will leverage cell transfer models or AM-specific gene knockout models to define how these pathways function *in vivo* to obtain a more holistic view of AM function.

In conclusion, this study shows that FLAMs are a useful model to interrogate mechanisms that make AMs unique among macrophage subsets. IFNγ responses are differentially regulated in AMs and BMDMs, and GSK3α/β modulates inflammation and T cell activation by AMs. Together, these findings build upon previous studies that suggest that there are key mechanistic differences between AMs and BMDMs and provide tools to better understand these differences and their roles in maintaining pulmonary function in health and disease.

## Materials and Methods

### Animal Experiments

All cell isolation involving live mice was performed in accordance with the recommendations from the Guide for the Care and Use of Laboratory Animals of the National Institutes of Health and the Office of Laboratory Animal Welfare. Mouse studies were performed using protocols approved by the Institutional Animal Care and Use Committee (IACUC). All mice were housed and bred under specific pathogen-free conditions and in accordance with Michigan State University (PROTO202200127) IACUC guidelines. All mice were monitored and weighed regularly. C57BL6/J mice (# 000664) and *Ifnar1-/-* mice (# 028288) were purchased from The Jackson Laboratory.

### Cell Isolation

J2 virus-immortalized Cas9^+^ BMDMs (iBMDMs) were isolated and immortalized from C57BL/6J mice as previously described (18, 56). FLAMs were isolated from C57BL/6J mice or *Ifnar1-/-* mice as previously described (21, 23). Briefly, fetal livers were extracted from euthanized dams immediately after sacrifice. Each liver was ground into a single-cell suspension and filtered through a 40-µm mesh screen and plated in one well of a six-well plate in FLAM media (see below). Primary AMs were isolated by bronchoalveolar lavage of C57BL/J6 mice, as previously described and cultured in FLAM media(57).

P25 TCR-Tg CD4^+^ T cells (40, 41) were isolated from the lymph nodes and spleens of transgenic P25TCR mice kindly shared by Joel Ernst. The spleen and lymph nodes were homogenized, passed through a 70-µm strainer, and washed with RPMI media (Gibco, Cat no. 11875093). CD4^+^ T cells were enriched using a MojoSort™ T cell isolation kit (BioLegend, Cat no. 480006) following the manufacturer’s protocol.

### Cell Culture

iBMDMs were maintained in Dulbecco’s Modified Eagle Medium (DMEM; HyClone Cytiva, Cat no. SH30243.01) supplemented with 10% fetal bovine serum (FBS) (R&D Systems, Cat no. S11550). iBMDMs were passaged once they reached 70–90% confluency. Cells were used in experiments after one week of culture. FLAMs were maintained in RPMI 1640 complete media supplemented with 10% FBS, 20 ng/mL recombinant human TGF-β1 (PeproTech, Cat no. 100-21), and 30 ng/mL recombinant murine GM-CSF (PeproTech, Cat no. 300-03). FLAM media was refreshed every 3 days. FLAMs were passaged at 70–90% confluency. Primary AMs were cultured in complete RPMI 1640 supplemented with 10% FBS, 30 ng/mL GM-CSF, and 20 ng/mL recombinant human TGF-β1. CD4^+^ T cells were cultured in complete RPMI 1640 media supplemented with 10% FBS and penicillin-streptomycin (50 U/mL penicillin, 50 mg/mL streptomycin) (Gibco, Cat no. 15140-122). All cells were incubated in 5% CO_2_ at 37°C.

### Macrophage Treatment Conditions

For all assays, unless otherwise indicated, FLAMs and iBMDMs were plated at a density of 1×10^6^ cells per well in 6-well tissue culture plates and were allowed to adhere overnight. The following day, cells were treated with DMSO (Fisher Chemical, Cat no. D128500), DMSO and 6.25 ng/mL IFNγ (PeproTech, Cat no. 315-05), 10 μM CHIR99021 (CHIR) (Sigma-Aldrich, Cat no. SML1046), or both 10 μM CHIR and 6.25 ng/mL IFNγ for 24 hours. In experiments with TNF (PeproTech, Cat no. 31501A) or IFNβ (BioLegend, Cat no. 581304), recombinant cytokines were added at 20 ng/mL for 24 hours. For the TNFR blocking experiment, an anti-TNFR blocking antibody (BioLegend, Cat no. 113104) was added at 1.25 ng/mL 24 hours prior to IFNγ activation.

### Flow Cytometry

For all experiments, cells were lifted by gentle scrapping, washed with PBS, and stained with MHCII-FITC (BioLegend, Cat no. 107606), CD40-APC (BioLegend, Cat no. 124612), PDL1-BV421 (BioLegend, Cat no. 124315), CD80-PE (BioLegend, Cat no. 104708), and CD86-APC-Cy7 (BioLegend, Cat no. 104708) TLR2-APC (Biolegend, Cat no. 153006) MRC1-PE (Biolegend, Cat no. 141706) Siglec1-FITC (Biolegend, Cat no. 142406) CD14-PE-Cy7 (Biolegend, Cat no. 123316) CD11a-PE (Biolegend, Cat no. 153103) (all diluted 1:400 in PBS). Cells were then washed 3 times in PBS and fixed with 1% formaldehyde (J.T. Baker, Cat no. JTB-2106-01) in PBS. Flow cytometry was performed on a BD LSR II or an Attune CytPix at the MSU Flow Cytometry Core, and data were analyzed using FlowJo (Version 10.8.1).

### Cytokine Profiling

FLAMs and iBMDMs were treated as described above for 24 hours, and supernatants were collected for cytokine profiling by Eve Technologies using a Mouse Cytokine/Chemokine 31-Plex Discovery Assay® Array.

### RNA Sequencing and Analysis

FLAMs and iBMDMs were plated in 6-well plates at 1×10^6^ cells/well and treated with IFNγ and CHIR as described above for 24 hours. The Direct-zol RNA Extraction Kit (Zymo Research, Cat no. R2072) was used to extract RNA according to the manufacturer’s protocol. Quality was assessed by the MSU Genomics Core using an Agilent 4200 TapeStation System. Libraries were prepared using an Illumina Stranded mRNA Library Prep kit (Illumina, Cat no. 20040534) with IDT for Illumina RNA Unique Dual Index adapters following the manufacturer’s recommendations, except that half-volume reactions were performed. Generated libraries were quantified and assessed for quality using a combination of Qubit™ dsDNA HS (ThermoFischer Scientific, Cat no. Q32851) and Agilent 4200 TapeStation HS DNA1000 assays (Agilent, Cat no. 5067-5584). The libraries were pooled in equimolar amounts, and the pooled library was quantified using an Invitrogen Collibri Quantification qPCR kit (Invitrogen, Cat no. A38524100). The pooled library was loaded onto 2 lanes of a NovaSeq S1 flow cell, and sequencing was performed in a 1×100 bp single-read format using a NovaSeq 6000 v1.5 100-cycle reagent kit (Illumina, Cat no. 20028316). Base calling was performed with Illumina Real Time Analysis (RTA; Version 3.4.4), and the output of RTA was demultiplexed and converted to the FastQ format with Illumina Bcl2fastq (Version 2.20.0).

All RNAseq analyses were performed using the MSU High Performance Computing Center (HPCC). Read quality was assessed using FastQC (Version 0.11.7) (58). Read mapping was performed against the GRCm39 mouse reference genome using Bowtie2 (Version 2.4.1)(59) with default settings. Aligned read counts were assessed using the FeatureCounts function from the Subread package (Version 2.0.0) (60). Differential gene expression analysis was conducted using the DESeq2 package (Version 1.36.0) (61) in R (Version 4.2.1). One IFNγ-treated FLAM sample did not pass QC and was not included in analysis. All raw sequencing data, raw read counts, and normalized read counts are available through the NCBI Gene Expression Omnibus (GSE239280).

For comparison to the Immgen dataset, core AM upregulated signature genes were compared between FLAMs and iBMDMs from our study and AMs and peritoneal macrophages from the Immgen Consortium (GSE122108). Raw counts were compiled and normalized in DESeq2. A box plot was generated in GraphPad Prism using normalized counts for core AM upregulated signature genes. GSEA analysis was used to identify enriched pathways in the RNAseq data set. Genes in the indicated comparisons were ranked using DeSeq2 and the “GSEA Preranked” function was used to complete functional enrichment using default settings for Hallmark Pathways from Mice. We acknowledge our use of the gene set enrichment analysis, GSEA software, and Molecular Signature Database (MSigDB) ((62), http://www.broad.mit.edu/gsea/).

### T Cell Assays

A previously established coculture system to assess antigen-specific T cell activation was used (63). In short, CD4^+^ T cells were stimulated with p25 peptide-pulsed (sequence: FQDAYNAAGGHNAVF from Genescript) iBMDMs or FLAMs that were irradiated with mitomycin (25 µg/mL) (VWR, Cat no. TCM2320) and had been pre-treated with DMSO, IFNγ, CHIR, and CHIR/IFNγ, as described above. Cocultures were supplemented with 10 ng/mL IL-12 (Peprotech, Cat no. 210-12) and 10 µg/mL anti-IL-4 (BioLegend, Cat no. 504-102) to achieve Th1 polarization. Supernatants were collected 3 days after coculture initiation and used to quantify T cell activation levels with ELISA (BioLegend, Cat no. 430801) following the manufacturer’s protocols.

## Acknowledgements

We thank Jessica F. Olive, PhD for helpful discussions and editing the manuscript. We thank members of the Olive lab for helpful discussions. The Attune CytPix, located in the MSU Flow Cytometry Core Facility, is supported by the Equipment Grants Program, award #2022-70410-38419, from the U.S. Department of Agriculture (USDA), National Institute of Food and Agriculture (NIFA). This work was funded by National Institutes of Health grants R35GM146795 and R01AI165618 to Andrew Olive.

---

**Table S1.** Normalized counts from all RNAseq samples.

**Table S2.** Ranked lists used for Gene Set Enrichment Analysis (GSEA)

**Table S3.** Normalized counts used for Immgen Comparative Analysis

**Table S4.** Normalized counts used for Heatmap comparisons

